# Cytonuclear analysis of barley spike traits using a cytoplasm-aware population

**DOI:** 10.1101/2025.04.08.647843

**Authors:** Schewach Bodenheimer, Eyal Bdolach, Avital Be’ery, Lalit Dev Tiwari, Ruth Sarahi Perez-Alfaro, Shengming Yang, Daniel Koenig, Eyal Fridman

## Abstract

The interplay between nuclear and cytoplasmic genomes—collectively known as cytonuclear interactions (CNIs)—is increasingly recognized as a key driver of phenotypic variation and adaptive potential across diverse organisms. Yet, leveraging cytoplasmic diversity and fully understanding CNIs’ contributions to agriculturally important traits remain major challenges in crop improvement, largely due to the scarcity of tailored genetic resources. In cultivated barley (*Hordeum vulgare*), limited genetic diversity relative to its wild relatives constrains adaptability to changing environments. While wild germplasm offers a reservoir of valuable alleles, the role of cytoplasmic variation and CNIs in shaping complex traits is still poorly understood. To address this gap, we present the Cytonuclear Multi-Parent Population (CMPP)—a novel interspecific resource comprising 951 BC_2_DH lines, generated from crosses between ten genetically diverse wild barley accessions (*H. vulgare* ssp. *spontaneum*) used as female founders and the elite cultivar Noga. This design facilitates the concurrent segregation and analysis of nuclear introgressions involving distinct wild versus cultivated cytoplasmic backgrounds from ten subfamilies. Phenotyping across multiple environments revealed that up to 5% of variation in key spike and grain trait BLUPs are explained by cytoplasm (η² = 0.05), including Thousand Grain Weight (TGW), Grain Width (GW), and Fruiting Efficiency at Maturity (FEm). Notably, wild cytoplasms influenced trait stability, with the B1K-50-04 cytoplasm increasing TGW stability based on Shukla’s measure. Genome-wide association studies (GWAS) employing Nested Association Mapping (NAM), FASTmrMLM, and MatrixEpistasis (ME) identified 76 marker-trait associations (MTAs). The ME approach specifically uncovered 16 cytonuclear QTL (cnQTL) exhibiting cytoplasm-dependent effects. Furthermore, we developed a genomic prediction (GP) strategy incorporating interactions between significant MTAs and population structure variables (subfamily and cytoplasm). This targeted interaction model (“Peaks + I”) achieved cross-validation accuracies comparable to, or even exceeding, models using the full set of 6,679 SNPs, despite utilizing substantially fewer predictors, enabling quicker and more efficient validation runs. The CMPP provides a unique platform for dissecting cytoplasmic effects and CNIs, highlighting the importance of incorporating cytonuclear context in genetic mapping and prediction to effectively harness both nuclear and cytoplasmic diversity for crop improvement.

## Introduction

Alongside the nuclear genome, cytoplasmic diversity has emerged as a significant adaptive driving force in plants (Bock, Andrew and Rieseberg, 2014). The interactions between nuclear and cytoplasmic DNA, known as cytonuclear interactions, have been shown to shape various traits across plant species (Atienza *et al*., 2008; Joseph *et al*., 2013; Roux *et al*., 2016) and in model organisms such as yeast (Nguyen *et al*., 2023). In barley, studies using chloroplast substitution lines have demonstrated the influence of cytoplasmic variation on important agricultural traits such as spike length, grain number, and grain weight (Goloenko *et al*., 2002). Moreover, recent research has revealed the impact of cytonuclear interactions on circadian traits and barley plant growth (Bdolach *et al*., 2019; Tiwari *et al*., 2024). The extent of variation explained by cytonuclear interactions varies across studies and traits. In *Arabidopsis thaliana*, cytonuclear epistasis was found to explain up to 17.8 % of the phenotypic variance for non-photochemical quenching (NPQ) and 12.3 % for projected leaf area (Flood *et al*., 2020). In yeast, Nguyen et al. (2020) reported that mitochondrial-nuclear interaction accounted for 10.8 to 31.5 % of phenotypic variation in colony growth, depending on the induced condition (Nguyen *et al*., 2020). Recent advancements in genome wide association study (GWAS) methodology, such as the SNPxGB and HBxGB models developed by Hamazaki et al. (2025), offer new tools for detecting quantitative trait loci (QTLs) that interact with complex population structures (Hamazaki, Iwata and Mary-Huard, 2025). Nevertheless, a systematic, mapping-based approach to localize specific cytoplasm-interacting nuclear regions underlying phenotypic variation in crops has been lacking. Although there are studies in other crops such as maize, where a biparental, reciprocal F_2:3_ population was used to identify cytonuclear epistatic loci affecting ear and plant height (Tang *et al*., 2013), their non-permanent nature precludes analysis of cytoplasm or cytonuclear interactions across different environments (GxE interactions).

Barley (*Hordeum vulgare*), a globally significant cereal reported as the highest production crop in Spain, Ireland, Finland, Norway, and Iceland in 2022 (*FAOSTAT*), provides an excellent system to explore these cytonuclear complexities in a major crop. Its versatility as a source of food, animal feed, and brewing material has cemented its importance in diverse agricultural systems worldwide. The remarkable adaptability of barley to a wide range of agroclimatic conditions stems from its rich genetic diversity, particularly evident in its wild relatives and landraces (Russell *et al*., 2016). On the other hand, cultivated barley varieties often suffer from a depleted allele pool, which poses a challenge to yield stability under climate fluctuations. Consequently, wild relatives have long been recognized as a potential source of beneficial alleles to enhance the vigor of established varieties (Warschefsky *et al*., 2014; Flint-Garcia *et al*., 2023). To effectively utilize these genetic resources, researchers have developed various population structures and mapping strategies. In 2008, Yu et al. introduced the first Nested Association Mapping (NAM) population in maize, which consisted of 5,000 recombinant inbred lines (RILs), derived from crosses between 25 diverse founders and a common parent (Yu *et al*., 2008). The NAM design is particularly advantageous because it incorporates a diverse pool of alleles while maintaining a structured format, and it combines the benefits of linkage mapping (requiring fewer markers) with the advantages of association mapping (offering high diversity and resolution) (Yu *et al*., 2008). In barley, three NAM populations have been published in recent years: The HEB-NAM (Maurer *et al*., 2015), AB-NAM (Nice *et al*., 2016), and BRIDG6 (Hemshrot *et al*., 2019). The HEB-NAM and AB-NAM both feature 25 wild or *agriocrithon* barley parents crossed back to a recurrent cultivated accession, while BRIDG6 contains the genomes of 88 cultivated accessions segregating in a cultivated background. These populations have facilitated the identification of quantitative trait loci (QTLs) for various traits ranging from flowering (Maurer *et al*., 2015), yield (Nice *et al*., 2017; Merchuk-Ovnat *et al*., 2018; Sharma *et al*., 2018), grain elements, (Herzig *et al*., 2019), pathogen resistance (Büttner *et al*., 2020, p. 20), and circadian output (Prusty *et al*., 2021). Spike and grain-specific traits, particularly TGW, have been the subject of extensive research due to their impact on yield. Hadjichristodoulou (1990) found that TGW was among the most stable yield-related traits across environments in barley, second only to plant height. The study also reported that TGW was positively correlated with grain yield, with correlation coefficients ranging from 0.32 to 0.40 in several trials (Hadjichristodoulou, 1990). Using a barley multi-parent population, Shrestha et al. (2022) found high (84 % or above) broad-sense heritability for grain size traits. Through comparing allelic effects between landraces and cultivars, they revealed that cultivars had a slightly higher proportion of alleles (54 % vs 46 %) contributing positively to grain size and weight, which suggests that larger and heavier grains have been favored in recent barley breeding efforts (Shrestha *et al*., 2022). Still, the study found that landraces also contributed significant positive alleles for grain size traits, indicating the potential value of wild relatives in breeding programs.

Besides linkage mapping and GWAS, genomic prediction has emerged as a powerful tool in plant breeding, allowing for the estimation of breeding values for complex traits using genome-wide marker data (Meuwissen, Hayes and Goddard, 2001). This approach has shown considerable promise in accelerating genetic gain in crop improvement programs by enabling early selection of superior genotypes and reducing the need for extensive phenotyping (Crossa *et al*., 2017). In barley, genomic prediction has been successfully applied to various traits, including plant height, leaf rust, malting and grain quality (Nielsen *et al*., 2016; Schmidt *et al*., 2016; Sweeney *et al*., 2020). With most prediction models calculating breeding values based on additive marker effects, recent studies have suggested that incorporating epistatic effects can improve prediction accuracy, when marker density and coding are chosen appropriately (Martini *et al*., 2017; Schrauf *et al*., 2020). Another recent study also suggested that modeling epistatic interactions between wheat-subgenomes could improve genomic selection and prediction accuracy (Tessele *et al*., 2025). These studies, however, are focused only on modeling interactions between nuclear SNPs, and sophisticated statistical techniques are needed to prevent high computational loads (Jiang and Reif, 2015).

In this study, we present a novel interspecific population called the Cytonuclear Multi-Parent Population (CMPP). The CMPP is designed to systematically investigate the interplay between nuclear and cytoplasmic genomes in barley, building upon the strengths of previous multi-parent populations while incorporating a deliberate focus on cytoplasmic variation obtained from different wild barley sources. We used the CMPP to investigate grain and spike-related traits in barley, including thousand grain weight (TGW), grain area (GA), grain length (GL), grain width (GW), length-to-width ratio (LWR), spike weight (SPW), grains per spike (GPS), fruiting efficiency at maturity (FEm), and grain weight per spike (GWPS). The CMPP allowed us to identify cytonuclear quantitative trait loci (cnQTL) controlling several of these traits. Moreover, we employed a multiplicative model that captures interactions between nuclear SNPs and population structure variables, such as subfamily and cytoplasm. This approach achieved cross-validation accuracy equal to or higher than that of models using all nuclear SNPs as predictors, while reducing computational cost. Our method applies to any multi-parent population, provided that there is sufficient statistical power, adequate group sizes, and sufficient frequency of each allelic combination.

## Materials and Methods

### Plant Material

The CMPP population includes nine wild barley accessions (*H*. *spontaneum*) from the Barley1K collection (Hübner *et al*., 2009) and one additional wild accession, HID386, from the IPK collection (Maurer *et al*., 2015). The *H. vulgare* cultivar Noga, a leading barley line in Israel, serves as the cultivated background. The wild accessions are from Yerucham (B1K-02-02), Michmoret (B1K-03-09), Ein Prat (B1K-04-04), Ashqelon (B1K-09-07), Mount Arbel (B1K-29-13), Mount Harif (B1K-33-09), Jordan Canal (B1K-42-16), Mount Eitan (B1K-49-18), Mount Hermon (B1K-50-04), and Kisalon (HID386) (see detailed geographic locations at (Hübner *et al*., 2009)).

Each of the 10 wild accessions was crossed as females with Noga to produce F_1_ hybrids. These F_1_ hybrids were backcrossed again as females with Noga to create BC_1_ plants. The 100 BC_1_ lines, subdivided to ten per wild parental origin, were then used for reciprocal backcrosses with Noga, resulting in 192 BC_2_ grains that served as starting point for doubled haploidization (DH, Figure 1A). The DH process was performed by SAATEN-UNION Biotec GmbH (Gatersleben, Germany). Sowing began in February 2020, with harvest from May to mid-July 2020. After a 3-month lab phase, the first regenerants were transferred to soil in August 2020. Seed setting occurred from late 2020 to January 2021. During the first half of 2021, the BC_2_DH CMPP lines were propagated in quarantine under natural conditions at ARO. The process resulted in 10 subfamilies, each divided into lines with either wild or cultivated cytoplasm (Figure 1B).

**Figure 1.**
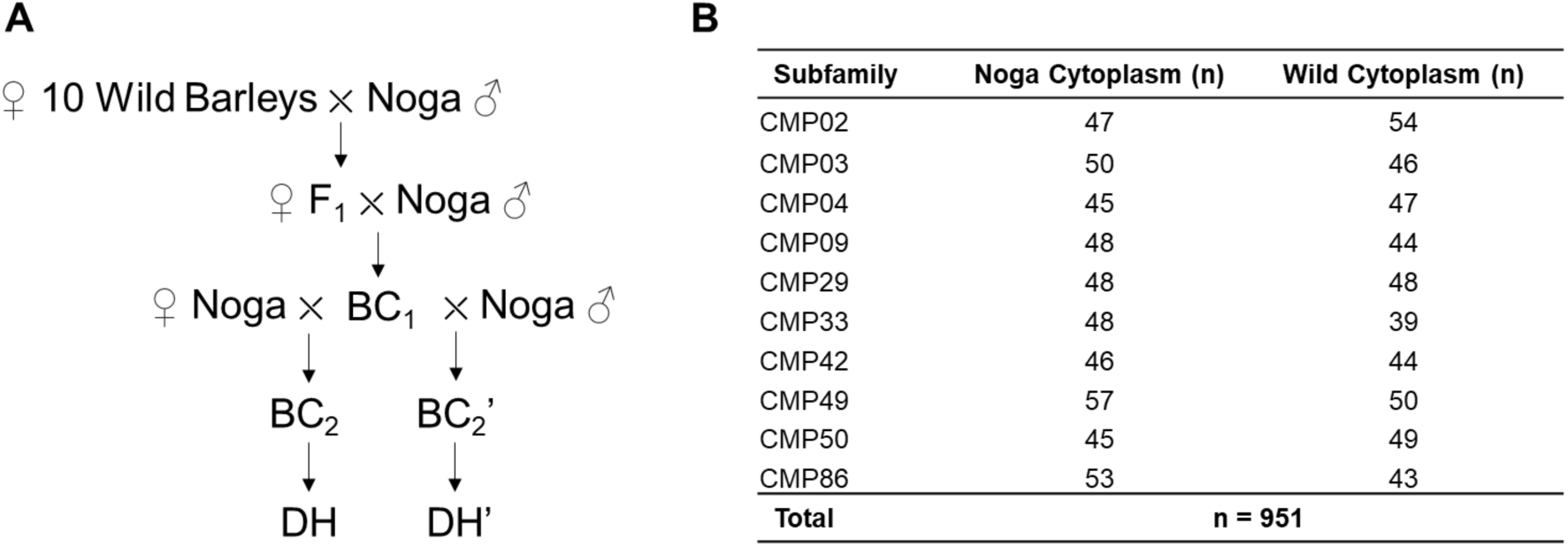
The interspecific barley cytonuclear multiparent population (CMPP). (A) Schematic representation of the crossing scheme used to generate segregation of nuclear and cytoplasmic genomes. The apostrophes at the BC_2_ and DH levels indicate different cytoplasms, resulting in subfamilies subdivided into wild or Noga cytoplasm carriers. (B) The CMPP population consists of 951 DH lines, each subdivided into subfamilies based on cytoplasmic background.

### Field Trials and Phenotyping

Field trials for the CMPP population were conducted at four locations: Rehovot, Yotveta, Yizream, and Mevo Hama. The trials in Rehovot and Yotveta were irrigated. The Rehovot trial occurred during the 2021/2022 season, while the trials at Yotveta, Yizream, and Mevo Hama were conducted during the 2022/2023 season. Trials were conducted without replication. In Rehovot, six plants were tray-germinated and grown in a greenhouse with 30 cm spacing between plots, and only TGW was measured. In the remaining sites, fifty grains from each line were divided evenly between two chambers in twelve-chamber magazines (Wintersteiger, Austria) and sown mechanically into two rows of 1 meter each, with 25 plants per row. Each line had two empty rows between them and a 1 m gap between plots. At full maturity, ten spikes were harvested from each plot, threshed, and used for all measurements. Grain size and counting were performed using a Marvin ProLine II seed analyzer (MARViTECH GmbH, Wittenburg). In total, the measured phenotypes can be subdivided into two classes: Grain-specific (TGW, Thousand Grain Weight; GW, Grain Width; GL, Grain Length; GA, Grain Area; LWR, Length-Width-Ratio), and spike-specific (SPW, Single Spike Weight; GWPS, Grain Weight per Spike). The difference between SPW and GWPS is that for GWPS, the chaff weight is not added to the weight of the spike, therefore representing the reproductive weight of the spike. FEm is calculated by dividing the number of grains per spike by the mature spike’s dry weight (Pretini *et al*., 2020).

### Whole genome sequencing

Lyophilized leaf tissue was disrupted for 964 DH progeny and the 11 population parents in 96 well plates using a Qiagen TissueLyser II instrument and use to extract using a standard CTAB protocol. Libraries were constructed using the Illumina DNA Prep Kit with reagents adjusted to a half-reactions. Whole genome sequencing was carried out on a NovaSeq 6000 instrument using the paired-end 150 bp read setting by the DNA Technologies Core at UC Davis, CA with a target sequencing depth of 10x for each of the CMPP parents and 1x for the progeny.

### SNP discovery in the CMPP parents

The CMPP wild parents were aligned to the *H. vulgare* Morex v.3 genome assembly (Mascher *et al*., 2021) using bwa v0.7.17 (Li and Durbin, 2009) and sorted using samtools 1.19.2 (Danecek *et al*., 2021). Duplicate reads were marked using the MarkDuplicates algorithm in gatk v.4.1.4.1 (Auwera and O’Connor, 2020). Variant calls were generated using the bcftools multivariant caller (Danecek *et al*., 2021) and filtered to include only biallelic SNP sites that have a quality greater than 100, no missing data, no heterozygous calls, a minimum per sample depth of 3, and at least 8 observations of the alternate allele.

### Genotyping the CMPP progeny

Sequencing reads from the progeny were aligned as described for the parents. The resulting bam alignment files were then processed to determine observed sites at each parental polymporphic site using python’s pysam module removing samples with no coverage. Sites were called as homozygous if greater than 90% of reads carried either the reference or alternate allele with other sites called as heterozygous. The resulting genotype calls were merged into a single vcf and used as input for beagle v.5.2 (Browning, Zhou and Browning, 2018) for missing data imputation using the parental genotypes as the reference dataset.

The imputed genotype dataset was further processed using SNPrelate (Zheng *et al*., 2012) to remove markers with a Minor allele frequency (MAF) below 0.1 and to prune nearby markers using an LD criterion on 0.2 within a sliding window of 0.5 Mbp, resulting in 6,679 genomic SNPs (Table S1). For the chromosome-wise MAF (Minor Allele Frequency)-density analysis, a separate genotype file was generated using a global MAF threshold of 0.05, and MAFs were then re-calculated within each family to prevent bias from markers that are fixed in certain families. The final MAF per marker was then determined as the average of its MAFs from within the subfamilies it segregates. Chloroplast DNA was obtained from pooled, wild cytoplasm-carrying lines from each of the CMPP subfamilies. Chloroplasts and their DNA extraction was done as described before (Tiwari *et al*., 2024), and Illumina sequencing performed by Genotypic Inc. (Bangalore, India). Clustering of chloroplast DNA was done using ETE 3 (Huerta-Cepas, Serra and Bork, 2016, p. 3), with MAFFT alignment of consensus sequences followed by tree building with PhyML and 100 rounds of bootstrapping.

### Phenotype Modeling

BLUP values (Table S2) were calculated using the lmer procedure (Bates *et al*., 2024) in R by defining trial site as fixed, and CMPP line as random effect. Broad-sense heritabilities (H2) according to Cullis (Cullis, Smith and Coombes, 2006) were extracted from the lmer objects using the h_cullis function from the agriutilities package (Aparicio, Bornhorst and CIAT, 2024). Shukla’s stability variance (Shukla, 1972) was calculated using raw TGW values from all four sites using the metan package (Olivoto, 2022). Meteorological data was obtained from the Israeli Meteorological Service (https://ims.gov.il/en/data_gov). Pairwise t-tests of cytoplasms within subfamilies for TGW BLUPs were performed using emmeans (Lenth *et al*., 2024).

### GWAS and Genomic Prediction

We employed three complementary mapping approaches to identify marker–trait associations (MTAs) in our multi-parent population: the Nested Association Mapping (NAM) model, FASTmrMLM, and MatrixEpistasis. In addition, genomic prediction was performed using a Bayesian whole-genome regression framework. For the NAM approach (Xavier *et al*., 2015), the phenotypic vector y was modeled as 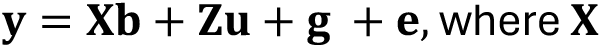 is the design matrix for fixed effects, **b** is the corresponding effect vector, **Z** is an incidence matrix linking individuals to subfamily-specific marker effects, **u** is the vector of these effects, **g** is the polygenic term and e represents the residual error. FASTmrMLM is an accelerated version of the mrMLM algorithm (Tamba and Zhang, 2018; Zhang *et al*., 2022). It is a biallelic model that does not consider subfamilies and uses a two-stage GWAS method, which first identifies candidate markers using a random-SNP-effect mixed linear model (RMLM), and then evaluates these markers collectively, which reduces the need for multiple testing correction, thereby reducing the type 1 error rate while maintaining statistical power. MatrixEpistasis was used to perform an exhaustive scan for epistatic interactions with full covariate adjustment, as outlined by Zhu and Fang (Zhu and Fang, 2018). The quantitative phenotype *p* is modeled as 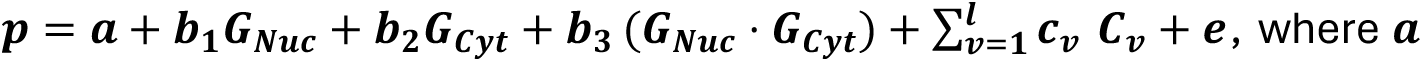 is the overall mean; ***G_Nuc_*** and ***G_Cyt_*** denote standardized values of nuclear SNPs and the cytoplasm-specific indicator variable, respectively; ***b_1_*** and ***b_2_*** are the additive effects; ***b_3_*** represents the interaction (epistasis) effect; *C**_v_*** (with ***v* = 1,…, 10**) are subfamily covariates with corresponding coefficients ***c_v_***; and ***e*** is the residual error.

In each method, BLUP values served as phenotypic inputs. FASTmrMLM directly provided final p-values, while p-values from NAM and ME were adjusted using chromosome-wise FDR correction. Significant MTAs were identified based on Bonferroni-adjusted p-values below 0.05. To determine the minimal distance between distinct MTAs, the average physical length corresponding to 5 centimorgan (cM) was calculated for each chromosome and used as a pruning criterion for nearby MTAs.

For genomic prediction, the bWGR implementation was used (Xavier, Muir and Rainey, 2020) which is based on a Bayesian whole-genome regression model 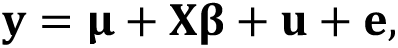 where **y** is the phenotypic vector, **μ** is the overall intercept, **X** is the design matrix containing marker data, **β** represents marker effects, and **u** is the polygenic term while **e** denotes residual error. The mcmcCV() function was used to simultaneously run the BayesA, BayesCPi, and BayesL models (Xavier, Muir and Rainey, 2020). To construct the genotype matrices used as input for prediction, a “peaks matrix” **M_Peaks_** was derived by subsetting the complete matrix **M** to include only the significant markers identified in the mapping analyses; thus, if ***H*** denotes the set of significant markers, then **M_Peaks_** = **M(:, H)**.

Additionally, a matrix with interactions was constructed by combining the interactive MTAs within **M_Peaks_** (**M_Subfam_** obtained from NAM mapping, and **M_Cyto_**obtained from ME mapping), with population structure variables. To that end, the design matrices **D_subfam_** and **D_Cyto_** were generated linking CMPP lines to their subfamily and cytoplasm, respectively. Using an element-wise product function denoted as **CME**, we then calculated the interaction matrices 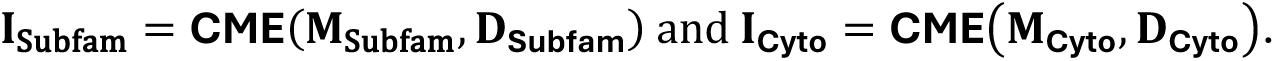 Finally, **M_Peaks+ I_** was obtained by merging **M_Peaks_**, **I_Subfam_**, and **I_Cyto_**. The cross-validation accuracy between **M_Peaks_**, **M_Peaks + I_**, and **M_full_** (the full genotypic matrix containing 6,679 SNPs) was then compared using 20 times fivefold, thus 100 cross-validations.

## Results

### Introgression analysis of the CMPP population

We conducted WGS-based genome-wide genotyping of the CMPP to serve as basis for genomic and association studies. In total, SNP-data was obtained for 940 lines, 920 of which were successfully phenotyped for at least a single trait. We then wanted to assess whether the within-family minor allele frequencies (MAFs) reflect the expected average introgression rate of 0.125 for BC_2_DH populations. The results showed that chromosome-wide MAFs range from 0.112 ± 0.03 (mean ± SD) for chromosome 4H, to 0.156 ± 0.049 for chromosome 3H (Figure 2, Table S3). The grand mean across all chromosomes is 0.137 ± 0.03, which is within 0.4 standard deviations of the expected 0.125.

**Figure 2.**
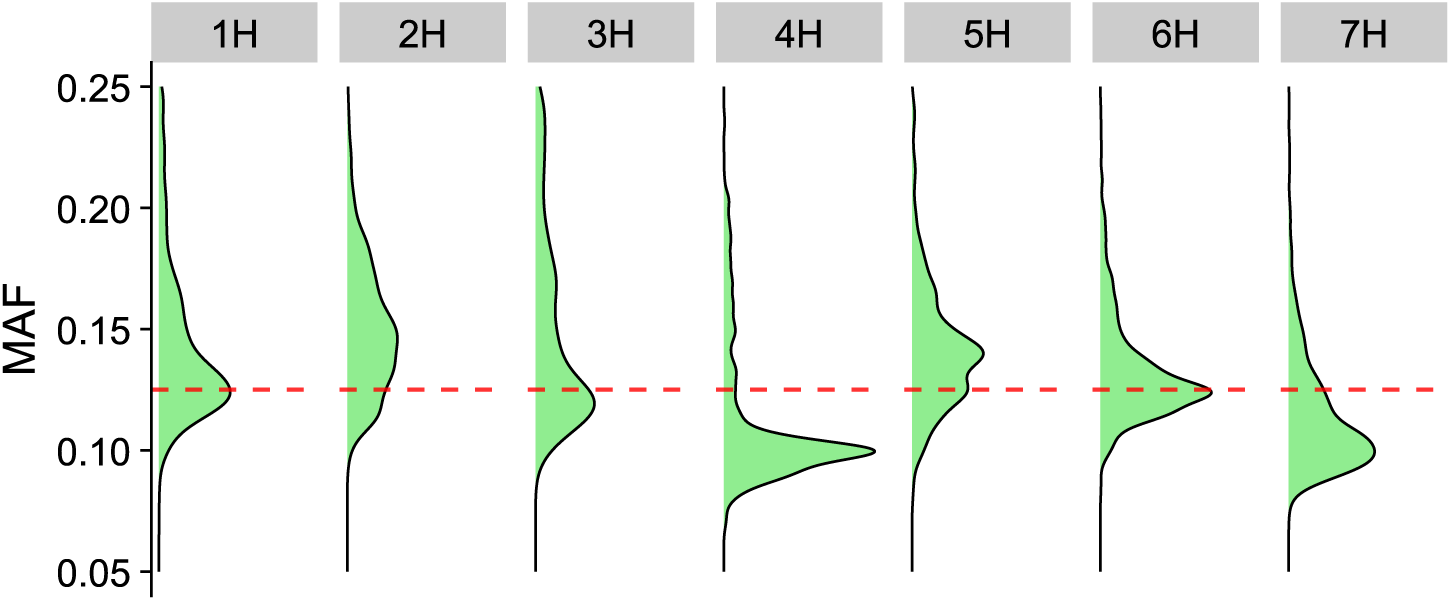
Density estimates of MAF for each chromosome in the CMPP, from markers pooled by family. The red line was placed at the expected 0.125 wild allele frequency for a BC_2_DH population.

### Quantifying Cytoplasmic Influence on Spike and Grain Phenotypes

We collected the phenotypes from field trials at four different ecogeographical zones and in a first step, wished to understand both the general trait heritability, as well as the contribution of subfamily and cytoplasm to overall variation. The linear models which therefore were fitted to the BLUP-values showed that GW had the lowest coefficient of variation (CV) at 2.56 %, whereas FEm had the highest at 7.42 % (Table 1). Generally, traits which we directly measured on the grains exhibited less variability than those we measured for the entire spike. This pattern was also reflected in the heritability estimates, where broad-sense heritability (H^2^) for TGW, GA, GL, GW, and LWR ranged between 0.65 and 0.69. In contrast, H^2^ for SPW, GPS, FEm, and GWPS ranged between 0.46 and 0.51.

**Table 1.** Summary statistics and heritability estimates for spike-related traits in the CMPP population. Traits measured include thousand grain weight (TGW), grain area (GA), grain length (GL), grain width (GW), length-to-width ratio (LWR), spike weight (SPW), grains per spike (GPS), fruiting efficiency at maturity (FEm), and grain weight per spike (GWPS). Values include mean, standard deviation (SD), coefficient of variation (CV), broad-sense heritability (H*^2^*), Eta squared (η^2^) and *R^2^* from two-way ANOVA. Envs represents the number of environments used to calculate the BLUPs. Significance: **** p≤0.0001, *** p≤0.001, ** p≤0.01, * p≤0.05, ns p>0.05.

| Trait | N | Envs | Mean | SD | Min | Max | CV (%) | $H^2$ | $\eta^2_{\text{Subfamily}}$ | $\eta^2_{\text{Cytoplasm}}$ | Total $\eta^2(R^2)$ |
| --- | --- | --- | --- | --- | --- | --- | --- | --- | --- | --- | --- |
| GA | 918 | 3 | 19.3 | 0.96 | 16.76 | 23.58 | 4.98 | 0.65 | 0.208**** | 0.020** | 0.228 |
| TGW | 951 | 4 | 44.18 | 2.73 | 34.67 | 52.59 | 6.17 | 0.68 | 0.170**** | 0.048**** | 0.218 |
| GW | 918 | 3 | 3.39 | 0.09 | 3.06 | 3.67 | 2.56 | 0.69 | 0.166**** | 0.050**** | 0.216 |
| GL | 918 | 3 | 8.14 | 0.35 | 7.23 | 9.48 | 4.26 | 0.66 | 0.157**** | 0.023** | 0.18 |
| LWR | 918 | 3 | 2.43 | 0.11 | 2.15 | 2.86 | 4.5 | 0.69 | 0.085**** | 0.044**** | 0.129 |
| GWPS | 903 | 3 | 1.19 | 0.09 | 0.78 | 1.46 | 7.42 | 0.47 | 0.092**** | 0.026** | 0.118 |
| SPW | 903 | 3 | 1.62 | 0.11 | 1.28 | 1.97 | 6.47 | 0.47 | 0.080**** | 0.029** | 0.109 |
| *FEm* | 898 | 3 | 67.91 | 8.25 | 43.55 | 97.51 | 12.15 | 0.46 | 0.053**** | 0.048**** | 0.101 |
| *GPS* | 904 | 3 | 26.89 | 1.71 | 17.61 | 32.39 | 6.37 | 0.51 | 0.045**** | 0.022* | 0.067 |

To test the contribution of cytoplasm to the overall BLUP variation for each trait, we fitted a two-way model and calculated the relative effect sizes as eta squares (η²) attributable to either subfamily or cytoplasm. As shown in Table 1, both subfamily and cytoplasm were significant factors for all traits. The highest η² values were noted for GW (0.05), TGW (0.048), and FEm (0.048), indicating that for these traits, around 5% of the total variation can be explained by the cytoplasm. Moreover, the variance explained by the subfamily was higher than the variance explained by cytoplasm for all measured traits. Examining the R² values of the two-way models, which also represent the combined η² for both subfamily and cytoplasm, the table reveals that GA had the highest combined variance explained (R² = 0.228), while GPS had the lowest (R² = 0.067), with grain-related traits again ranked at the top.

Building on these findings, we aimed to further distinguish the effects of cytoplasmic variation from those influenced by nuclear genetic differences within the CMPP. This distinction is important to ensure that any observed differences between groups with different cytoplasmic backgrounds are not just due to genetic drift during the CMPP’s breeding and development phase (Figure 1A). To do this, we calculated the relative effect of the wild cytoplasm (REW) and its significance for each trait and subfamily. We also used Principal Component Analysis (PCA) to visualize the population structures specific to each subfamily. Figure 3A illustrates that significant effects between cytoplasmic groups existed for all traits except for GA, GPS, and GWPS. The largest significant REW were observed for FEm in CMP29 (REW =-10.9 %, *p* < 0.001), SPW in CMP42 (REW =-5.4, *p* < 0.001), TGW in CMP29 (REW =-3.8%, *p* < 0.01), and LWR in CMP50 (-3.7%, *p* < 0.001). The trait with the most significant pairwise difference is GW (*n* = 4), while the subfamily with the most significant pairwise trait differences is CMP50 (*n* = 4). Furthermore, Figure 3B shows that the SNP-based clusters corresponding to different cytoplasmic groups within these subfamilies do not show divergence between cytoplasmic groups, supporting the conclusion that the observed cytoplasmic effects (Figure 3A) are genuine and not confounded by nuclear genetic drift.

**Figure 3.**
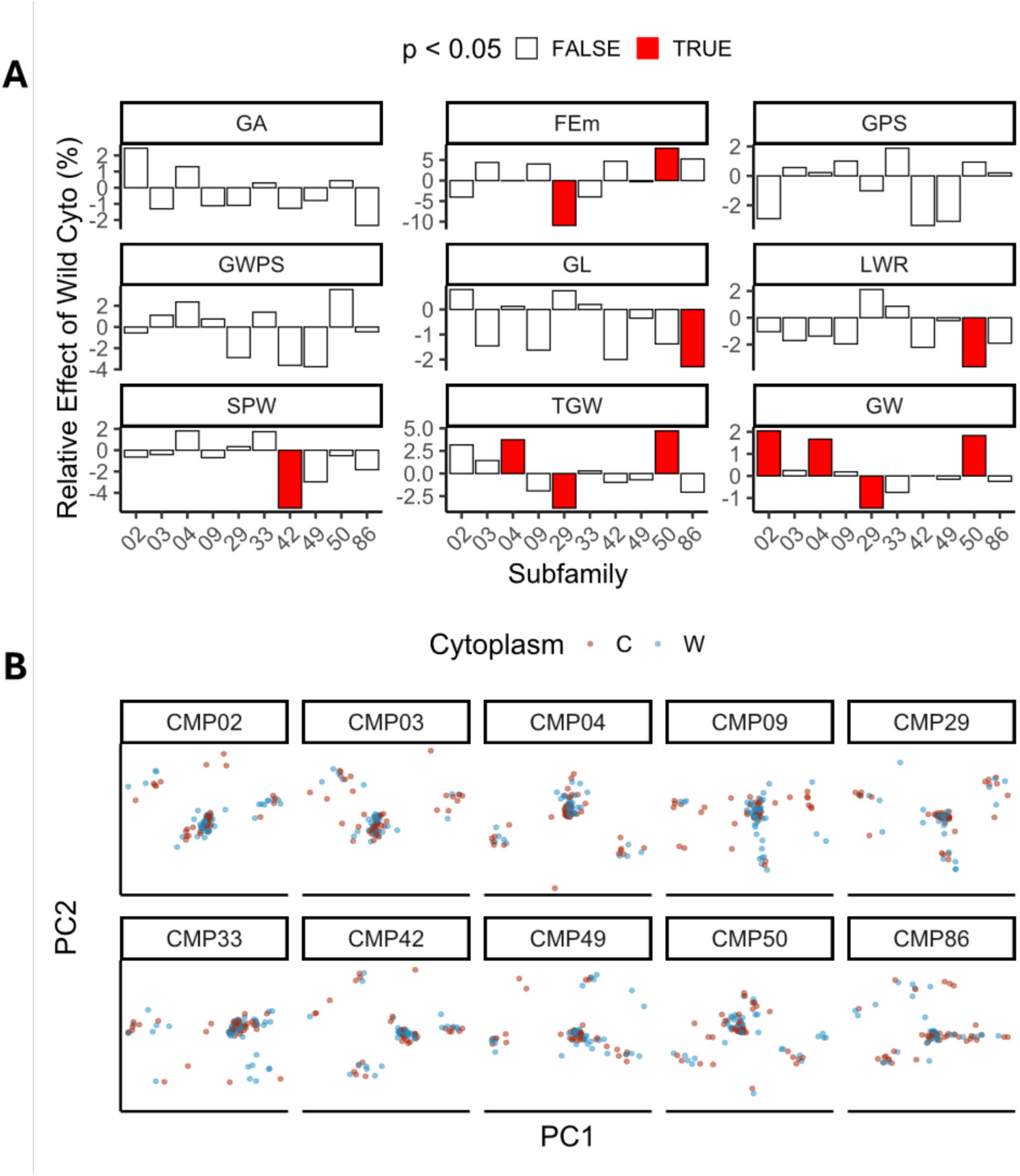
Analysis of cytoplasmic effects within subfamilies. (A) Relative effects of cytoplasm within each subfamily, for each trait. Bars show the relative effect of the wild cytoplasm-carrying group within the subfamily, with significant T-tests between cytoplasmic groups highlighted in red. C, Cultivated; W, Wild. (B) PCA of SNP data for each subfamily, color-coded by cytoplasmic group.

### Cytoplasmic Effects on Grain Weight Stability

Notably, the field trials were performed in varying environmental conditions ranging from Rehovot in Central Israel with two days of extreme heat stress during the growing season (>35°C) to Mevo Hama having 13 days of extreme heat stress (Figure 4A). We therefore wished to examine how trait stability is influenced by cytoplasm. To this end, we divided the CMPP into 20 groups, each representing a unique combination of subfamily and cytoplasm (either from Noga or the wild parent). For each group, Shukla’s stability variance parameter was calculated (Shukla, 1972). This parameter gives insight about the contribution of each genotype to the overall genotype-environment interaction, with lower values indicating more stable genotypes. While CMP02 and CMP50 both appear as outliers, the portion of CMP50 carrying the wild cytoplasm leads to rescue of stability (from 8.89 for the Noga, to 1.04 for the Wild cytoplasm), suggesting specific influences of certain cytoplasms on TGW robustness (Figure 4B).

**Figure 4.**
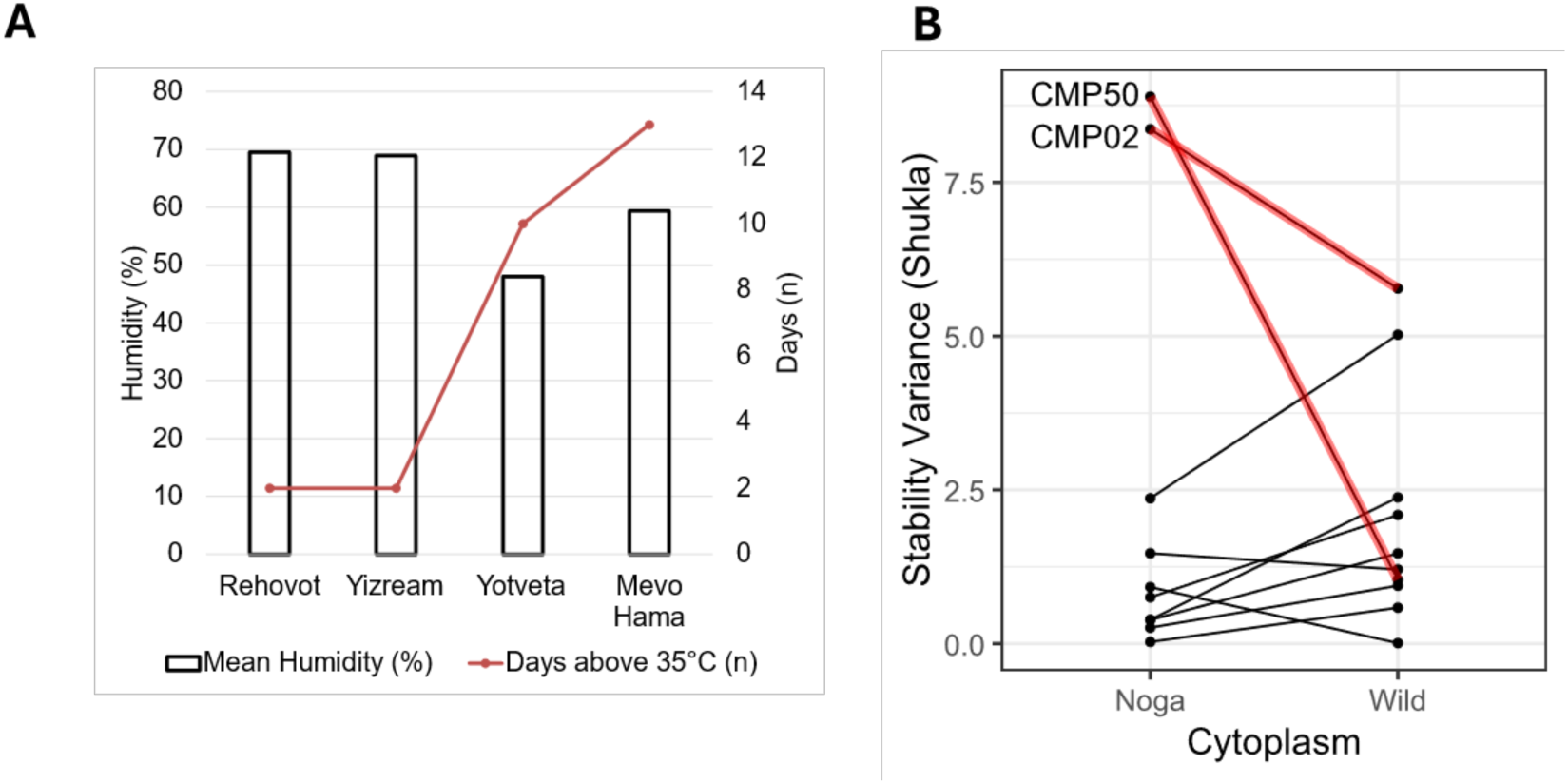
Influence of cytoplasm diversity on phenotypic stability across environments. (A) Total Precipitation, and number of days where the maximal temperature exceeded 35°C, as measured during the growing periods. (B) Shukla Stability Variance for both the Noga and Wild cytoplasm portions of the 10 CMPP subfamilies. Outliers CMP02 and CMP50 are marked for reference.

### Genome-wide association and cytonuclear interaction analysis

We used three methods to search for marker-trait associations (MTAs): the nested association mapping (NAM) model (Xavier *et al*., 2015)), the FAST multi-locus random SNP effect mixed linear model (FASTmrMLM, Wang *et al*., 2016; Li *et al*., 2022) and the MatrixEpistasis (ME) approach (Zhu and Fang, 2018). The NAM model, designed for NAM populations, allows the detection of subfamily-specific MTAs by varying marker coefficients according to the subfamily of origin. The FASTmrMLM model, which does not take subfamily indicators into account, increases the power to detect significant associations by including multiple markers simultaneously. The ME approach efficiently searches for interactions between pairs of markers while using predefined covariates to account for subfamilies, enabling the identification of cytonuclear QTL (cnQTL).

Overall, we identified 76 MTAs: 11 from the NAM model, 49 from FASTmrMLM, and 16 from ME (Figure 5, Table S4). As can be seen in Figure 5 and Table S4, the region between wgs33905 (335 Mbp) and wgs67051 (516 Mbp) on chromosome 1 features a notable cluster of 20 MTAs encompassing all traits except for GW, with GA, LWR, and GL all being represented by all three mapping methods. Chromosome 1 also harbors the highest number of trait-wide NAM-MTAs (n = 7), with other chromosomes featuring at most a single NAM-MTA trait-wide (Figure 5, Table S4).

**Figure 5.**
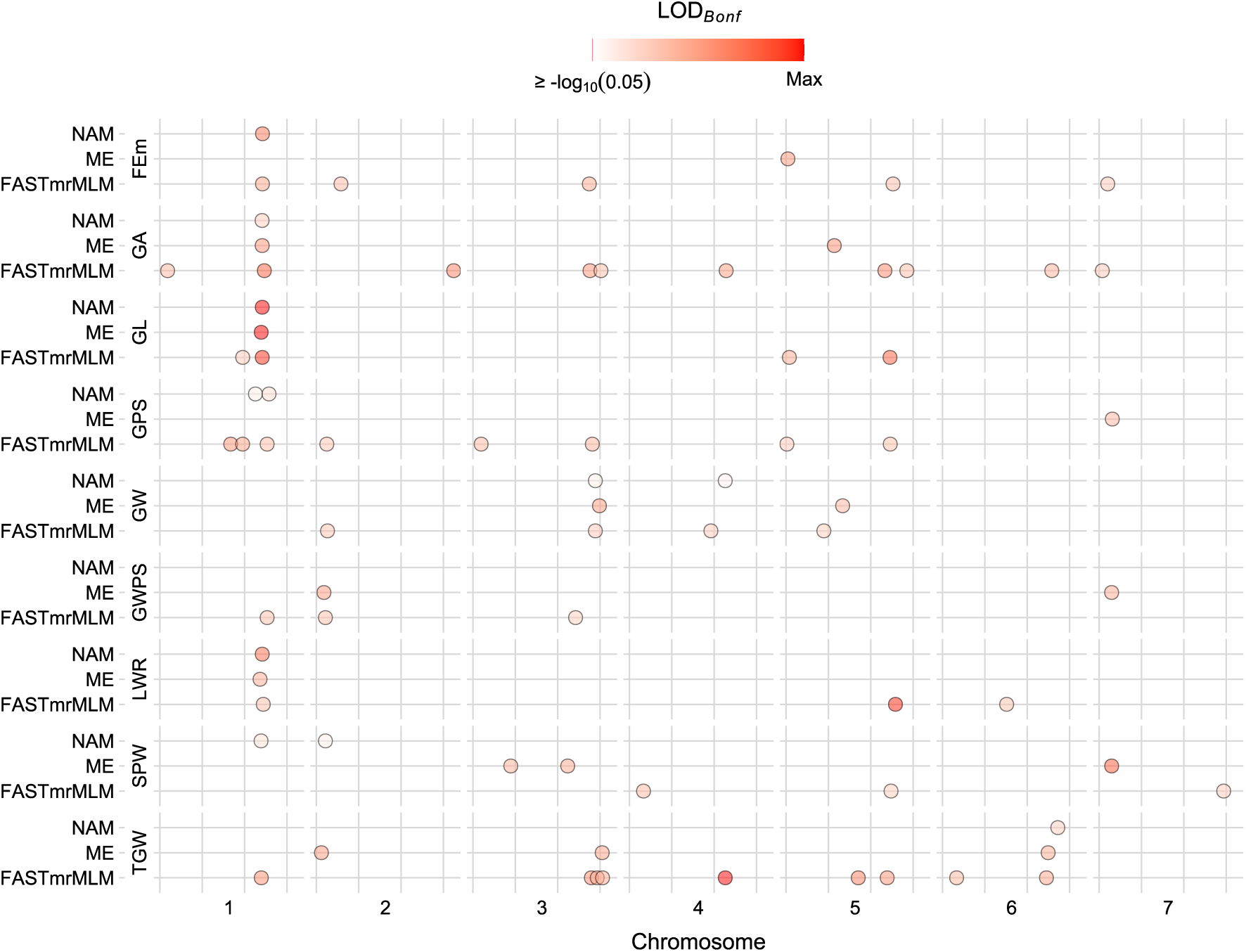
Chromosome-wise marker-trait associations (MTAs) identified using NAM, FASTmrMLM, and ME models. MTAs from all three methods are highlighted for each chromosome, with a color scale indicating the significance level of the association, scaled by the mapping method. NAM, Nested association mapping; FASTmrMLM, FAST multi-locus random SNP effect mixed linear model; ME, MatrixEpistasis.

The NAM approach’s ability to measure subfamily-specific effects is exemplified in Figure 6A, which shows the SNP effects on GL along chromosome 1H for subfamilies CMP02 and CMP04. Here, wild introgressions within CMP04 around the highlighted peak MTA *wgs58876* exhibit positive effects on GL, while wild introgressions within CMP02 are associated with negative effects. Figure S3 shows the GL effect profiles for all subfamilies, whereas CMP02 and CMP03 feature the strongest negative, and CMP04 and CMP29 have the strongest positive effect around *wgs58876*.

**Figure 6.**
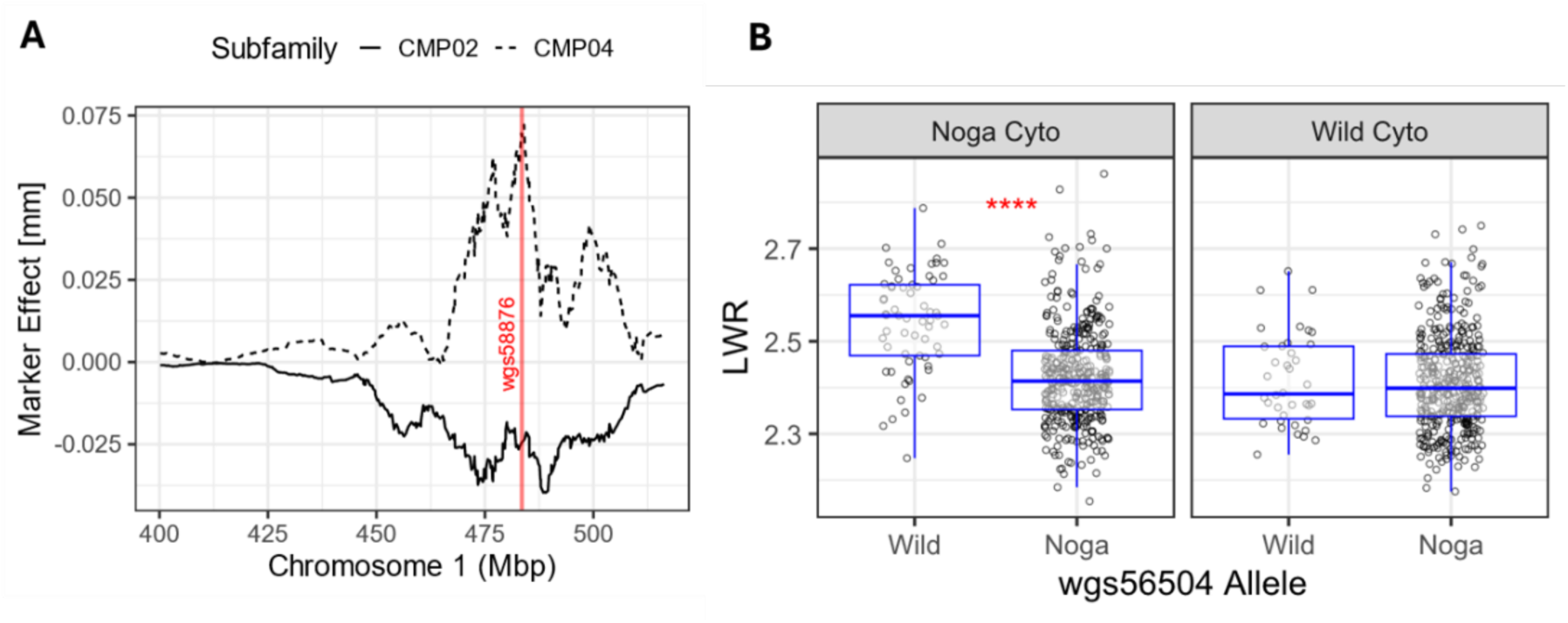
Subfamily-specific SNP effects on GL, as well as a cytonuclear interaction affecting grain length-to-width ratio (LWR) on chromosome 1H. (A) GL SNP effects along chromosome 1H for subfamilies CMP02 and CMP04, as obtained through the NAM procedure. Red line is positioned at the local peak *wgs58876*. The line represents the effect’s rolling mean with a window size of n = 20 markers. (B) Interaction between nuclear allele and cytoplasm at the LWR-cnQTL *wgs56504*. The marker is seen segregating in both cytoplasmic backgrounds. Significance: **** p≤0.0001, *** p≤0.001, ** p≤0.01, * p≤0.05, ns p>0.05.

Single-marker association testing of the cnQTL using ME demonstrates how the cytoplasmic background conditions allele effects at these loci. At the LWR-cnQTL *wgs56504* on chromosome 1H, the wild nuclear allele significantly increases the LWR in the Noga cytoplasmic background by 5%, while no effect is measured for *wgs56504* when the wild allele segregates in wild cytoplasmic background (Figure 6B). Figure S4 shows pairwise plots for all cnQTL detected in the scan. Strikingly, *wgs209080*, *wgs238755*, and *wgs76388* stand out by showing significant effects in both opposite cytoplasms for their respective traits (SPW, TGW, and GPS).

### Inclusion of cytonuclear interactions in genomic prediction

We next assessed the potential benefits of incorporating cytonuclear interactions into genomic predictions for TGW, SPW, GPS, and FEm. Specifically, we aimed to determine whether accounting for cytoplasmic relationships would enable us to capture additional genetic variation without incurring the complexity and computational demands typically associated with modeling epistatic interactions (Varona *et al*., 2018). To address this, we compared cross-validation accuracies obtained from three distinct marker matrices: (1) a baseline genotype matrix comprising all 6,679 SNPs (“All Markers”); (2) a reduced matrix (“Peaks”) containing only the markers from significant marker-trait associations (MTAs) identified by FASTmrMLM, NAM, and ME methods; and (3) an enhanced matrix (“Peaks + I”) that includes the significant markers from matrix (2), along with additional terms capturing interactions between NAM-peaks and subfamily genetic backgrounds, as well as between cnQTL (i.e., ME-peaks) and cytoplasmic backgrounds. Figure 7 illustrates the predictive accuracy achieved with these marker matrices using the three most effective genomic prediction models (BayesA, BayesCpi, and BayesL). Notably, for traits FEm, GPS, and TGW, the “Peaks + I” matrix achieved prediction accuracies equivalent to those of the “All Markers” matrix, despite utilizing only a fraction of the number of predictor variables (0.6%, 0.8%, and 1.2%, respectively). Notably, “Peaks + I” even significantly outperformed “All Markers” for BayesCpi in GPS, leading to a 11.2% relative increase in predictive accuracy (*p* = 0.047, Table S5).

**Figure 7.**
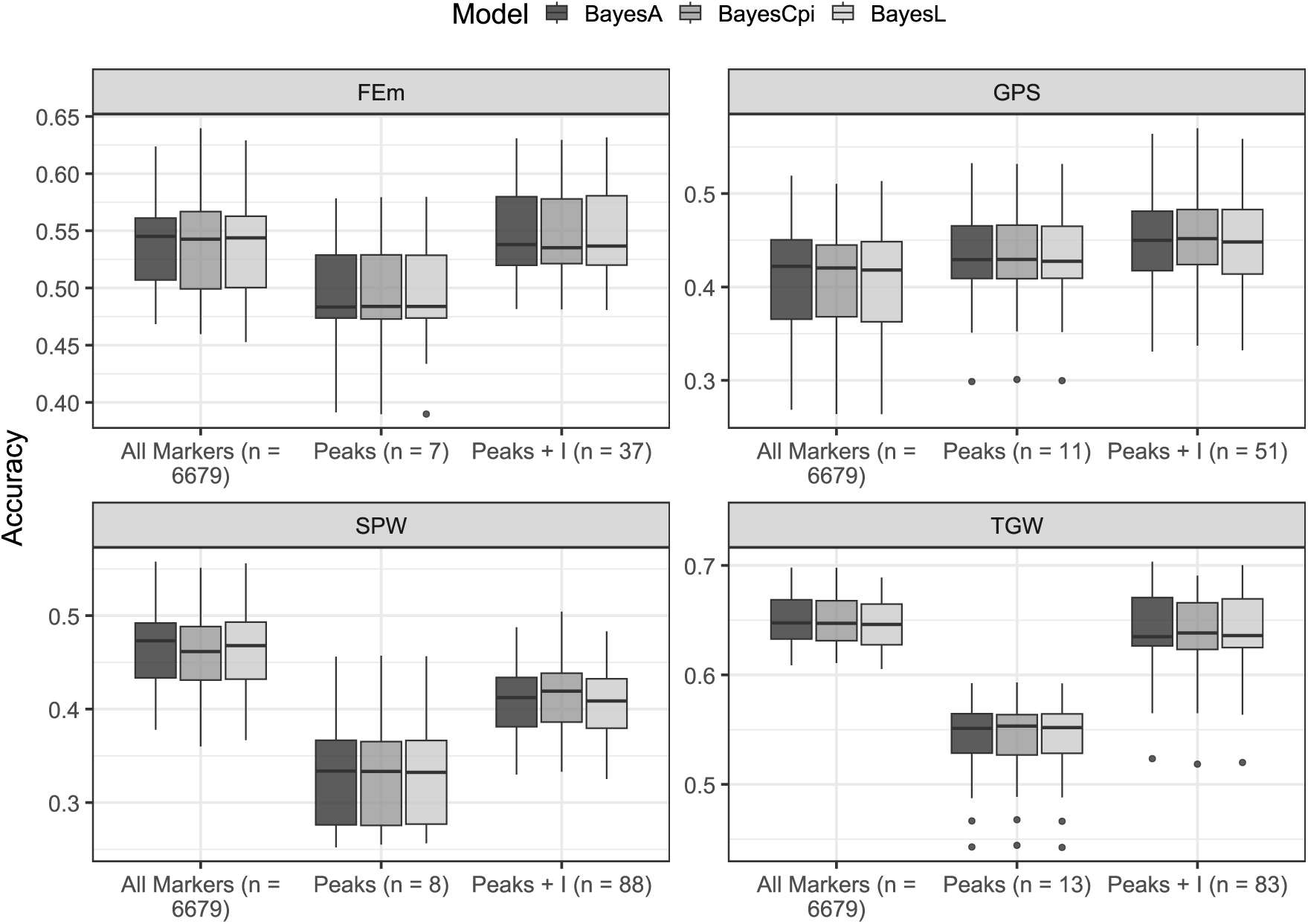
Comparative visualization of cross-validation accuracies across three predictor matrices in genomic prediction models. “All Markers”: Full genotypic matrix; “Peaks”: Pruned set of significant MTAs from all three mapping methods; “Peaks + I” All MTAs from “Peaks” combined with interaction terms obtained from peak markers and population structure indicators for thousand grain weight (TGW), spike weight (SPW), grains per spike (GPS), and fruiting efficiency at maturity (FEm).

## Discussion

### The CMPP as a New Genetic Infrastructure

As a new member of the barley multi-parent population (MPP) resources, the CMPP enables the linkage of untapped cytoplasmic variation from the wild for key agronomic traits. While previous barley MPPs have led to the successful identification of QTL controlling a number of yield-related traits (Maurer, Draba and Pillen, 2016; Merchuk-Ovnat *et al*., 2018; Afsharyan *et al*., 2023), the search for epistatic interactions did not consider parental effects, but was limited to epistasis between nuclear loci (von Korff, Léon and Pillen, 2010; Mathew *et al*., 2018; Afsharyan *et al*., 2020). While the DH character of the CMPP lines makes them an immortal mapping population with the possibility to validate QTL in different environments, the DH process was initiated directly on the BC_2_ plants before any inbreeding, thus preserving maximum genetic diversity among the lines. The CMPP also serves as an important addition to the preservation and utilization of allelic diversity of barley, by including several Barley1K accessions (Hübner *et al*., 2009) that originated from habitats further south in the Levant, compared to the lines included in the HEB population (Maurer *et al*., 2015). The most southern accession in the HEB population is HID386, which although it is also originated from Israel, as the other Barley1K accessions, it was collected in Kisalon (31.7743° N, 35.0482° E). This is compared to the inclusion of accession from Yerucham (B1K-02-02; 30.9872° N, 34.9310° E) and Mount Harif (B1K-33-09; 30.4936° N, 34.5608° E) within the CMPP (Fig. 1). The two latter accessions represent the more xeric parts of the Southern Levant (Chang *et al*., 2022) and therefore could provide relevant cytonuclear alleles conferring adaptation to more harsh environments, e.g. stabilizing effects of the CMP02 cytoplasm (Fig. 4B), which are absent in current MPP. This additional allelic repertoire would be imperative for gene discovery and for breeding better adapted barley for future scenarios of climatic change (Zhang and Batley, 2020).

### Chloroplast Diversity in the CMPP and implications to GWAS

We used two principal approaches to investigate cytoplasmic effects within the CMPP. First, as shown in Figure 3B and Figure 4A, we distinguished each CMPP subfamily by comparing cultivated and wild cytoplasmic counterparts. For ME mapping, on the other hand, cytoplasms were broadly categorized as ‘cultivated’ or ‘wild’. While this method disregards the individual nature of each wild cytoplasm, it was an essential choice to avoid sample size shrinkage which is a natural consequence of analyzing rare allelic combinations, thereby ensuring sufficient statistical power during interaction testing. Despite the broad categorization, we conducted whole-genome sequencing on the chloroplast sequences of each subfamily, revealing non-synonymous mutations at 12 chloroplast genes (Table S6), including variation at the RpoC1 gene locus, which we showed previously to influence the circadian clock plasticity (Tiwari *et al*., 2024). Importantly, a phylogram resulting from the consensus sequences from Noga and all of the wild chloroplast sequences, and including the *Hordeum vulgare* L. var. *trifurcatum* (Ren *et al*., 2021) chloroplast sequence as an outgroup (Figure 8), showed that Noga clusters distinctly from the wild barley chloroplast sequences, thereby providing support for using a binary cytoplasm classification for cytonuclear epistasis mapping. Nevertheless, future studies should also consider mitochondrial variation, although limited diversity between wild and cultivated barley mitochondrial sequences was found in such comparisons (Hisano *et al*., 2016).

**Figure 8.**
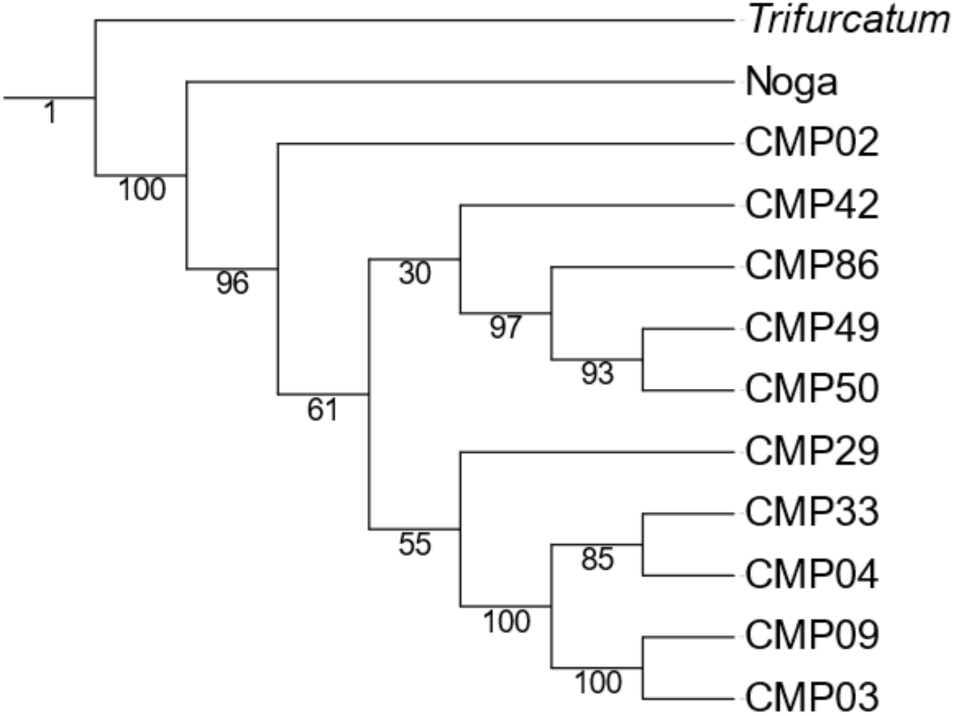
Maximum-Likelihood tree of chloroplast sequences. Clustering is based on a multiple sequence alignment of chloroplast sequences originating from Noga and the wild cytoplasms. *Hordeum vulgare* L. var. *trifurcatum* was used as an outgroup. Branch supports were computed out of 100 bootstrapped trees.

### Standing variation in the CMPP for spike and grain phenotypes

In this study, we focused on spike and grain traits to make the first attempt at dissecting standing variation using this new resource, primarily because these traits are known for their relatively high heritability (Sadras and Slafer, 2012). However, less is known about potential cytonuclear effects on these traits. The heritability of the spike-related traits in our study showed values consistent with those reported in other studies (Table 1). Importantly, genetic PCA of cytoplasmic groups within CMPP subfamilies confirmed the absence of population structure within subfamilies, strengthening the evidence for a cytoplasmic origin of the observed effects (Figure 3).

Fruiting efficiency at maturity (FEm), which is the proportion of chaff dry weight to the number of grains per spike, has shown a high narrow-sense heritability in wheat, making it a promising trait for indirect selection to improve grain number and yield (Alonso *et al*., 2018; Pretini *et al*., 2020). Although fruiting efficiency (FE), where spike dry weight is measured at anthesis, is a better predictor for final grain number, FEm remains a valuable proxy for breeding programs due to its simpler measurement compared to FE, especially in early generations with a high number of lines to evaluate (Pretini *et al*., 2020). Our investigation highlights the relevance of Fruiting Efficiency at Maturity (FEm) as a target trait in barley. Within the CMPP, FEm displayed moderate broad-sense heritability (H²=0.46, Table 1) and, critically, showed one of the strongest cytoplasmic influences (η²=0.048) among all spike and grain traits studied, comparable only to TGW and GW. This significant cytoplasmic effect, further evidenced by specific strong REW in subfamilies CMP29 and CMP50 (Figure 3A), validates the utility of the CMPP design for dissecting cytoplasmic control over this yield-related trait.

GWAS using three different approaches revealed method-specific MTAs that were either general, subfamily-specific or cytoplasm-conditional. Recently, Du et al. (2024) reviewed 54 papers and identified 85 meta-QTLs (MQTL) influencing barley yield and yield-related traits (Du *et al*., 2024). We compared the MTAs from all our GWAS analyses to these MQTL ranges and found that 48 out of the 76 totally identified MTAs fall within one or more MQTL-ranges (Table S4), with MQTL1H-8 (chromosome 1H, 435.27 - 502.06 Mbp) standing out with 14 MTAs identified in this study within its boundaries. Notably, the upper limit of MQTL1H-8 lies only 13 Mbp downstream from *<u>HvELF3</u>* (HORVU.MOREX.r3.1HG0095050). Variation at this gene was shown to impact traits ranging from photoperiodic flowering to plant development (Boden *et al*., 2014; Zahn *et al*., 2023), with some *HvELF3* mutations associated with the geographic expansion of barley (Faure *et al*., 2012; Zakhrabekova *et al*., 2012). When considering the value of scanning for cnQTL, an important question that arises is whether these loci would have been identified through more conventional mapping functions. As can be read from Table S4, 15 out of the 16 cnQTL lie no more than 40 Mbp away from an MTA identified through either NAM or FASTmrMLM, with wgs185810 (SPW) being an outlier located 140 Mbp away from the closest non-cytonuclear MTA. However, it is important to note that although some cnQTL are found in the vicinity of MTAs from other methods, this doesn’t necessarily mean that the same traits are influenced by these neighboring MTAs, which again suggests that using cytonuclear trait mapping could help to identify untapped variation.

To validate cnQTL and facilitate their application in breeding and gene discovery, it is necessary to aim for sufficiently high nuclear allele frequencies in selected cytoplasmic backgrounds. These follow-up experiments could adopt the family-based analysis in which selected CMPP lines carrying a phenotypically noticeable cytonuclear combination will be crossed as females and males, followed by selfing and comparative genotype-to-phenotype analysis between the reciprocal F_2_ populations. This is an approach that is currently being pursued through the development of reciprocal F_2_ populations, where the same candidate cnQTL segregate in contrasting cytoplasmic backgrounds thereby creating “quartet” of the four genotypic groups for comparison of the cytoplasm-conditioned effects of the nuclear QTL.

### Stabilizing Effects by B1K-50-04 Cytoplasm in Different Populations

The need for more resilient crops is a major driver of crop wild relatives research (Warschefsky *et al*., 2014; Hufford, Teran and Gepts, 2019; Toulotte *et al*., 2022). This study demonstrates that for the current phenotypic space there are at least two of the ten subfamilies exhibit a clear distinction between cultivated and wild cytoplasms regarding trait stability, including CMP50, which originated from a cross between Noga and B1K-50-04 (Hübner *et al*., 2009). In our case, and as indicated by Shukla’s stability variance, this cytoplasmic background restores trait stability within the CMP50 subfamily (Figure 4B). It has also been shown to confer higher thermal robustness of both circadian traits (Bdolach *et al*., 2019) and whole plant growth (Tiwari *et al*., 2024) in a wild barley reciprocal mapping population (Bdolach *et al*., 2019). In the same whole-wild population, derived from a reciprocal cross between two wild barleys, the B1K-50-04 cytoplasm was associated with inferior reproductive fitness, as indicated by SPW (Tiwari *et al*., 2024). Here, for the cultivated-wild interspecific CMPP, since SPW was not measured in four environments, which we considered the minimum for calculating robustness measures, we could not determine whether the robustness of TGW driven by wild cytoplasm in CMP50 persisted for SPW. This is especially relevant as SPW was more dependent on GPS than on TGW (r = 0.63 vs. r = 0.46, respectively, Figure S1), indicating a commonly observed trade-off in cereal crops (Quintero *et al*., 2018).

While insights into crop domestication are to be gained, it is important to consider that wild genetic elements enhancing the robustness of agricultural traits, such as the B1K-50-04 cytoplasm, may lead to pleiotropic effects. For example, during a study with an interspecific MPP, the Halle Exotic Barley (HEB; Maurer *et al*., 2015), the *DOC3.2* QTL on chromosome 3H increased biomass under higher temperatures but simultaneously reduced yield (Prusty *et al*., 2021). Nonetheless, we envision that the advanced nature of the CMPP will facilitate the utilization of wild nuclear and cytoplasm diversity in barley breeding.

### Implications of cnQTL and cytonuclear genomic prediction for breeding

To overcome the limitation of GP models that rely solely on additive marker effects, models such as Epistatic Random Regression BLUP (ERRBLUP) and selective Epistatic Random Regression BLUP (sERRBLUP) were developed (Vojgani *et al*., 2021). While the benefit of SNP-SNP interactions in GP has been put into question when marker density was high, epistatic models did outperform additive ones when a low marker density was chosen (Schrauf *et al*., 2020). Here, our GP model comes into play.

Our GP model introduces an innovative approach by incorporating cytonuclear interactions, providing a targeted and efficient alternative to traditional epistatic models. Instead of modeling all possible pairwise SNP interactions—which can be computationally intensive and prone to overfitting—our method specifically targets interactions between nuclear markers and key population structure variables: subfamily and cytoplasmic background. This strategy leverages the biological significance of cytonuclear interactions, validated by our mapping results, effectively capturing the subset of epistatic effects most influential to the traits of interest. As demonstrated with FEm, GPS, and TWG, incorporating these targeted interactions achieved cross-validation accuracies comparable to models utilizing the entire set of 6,679 genomic markers.

Nevertheless, this method is not without limitations, most notably the constraint imposed by the uniparental inheritance of cytoplasmic genomes. Unlike nuclear markers, which can be recombined and stacked through breeding cycles to accumulate favorable alleles, different cytoplasms cannot be combined within a single individual. While this inherent limitation reduces the flexibility of recurrent cytonuclear-based selection, we believe that (1) this method serves as an efficient tool for initial selection rounds, especially if comprehensive genotyping is not accessible, (2) the method enhances our understanding of marker effects by modeling how they vary within the context of cytoplasmic background and (3) it opens the door to re-evaluate data from existing structured plant populations where the directionality of crosses are known.

## Data Availability Statement

The authors affirm that all data necessary for confirming the conclusions of the article are present within the article, figures, and tables. The filtered VCF file including 556,286 sites was uploaded on Figshare. A self-contained code folder named “Supplementary Code” was also added in Figshare, which includes code and data to replicate the analyses.

## Acknowledgments

We would like to thank Arik Hirshman and his team in HaZera39 seeds for their help in sowing, growing the CMPP in different environments and for the availability of the Marvin machine for phenotyping. We also thank Booky Katz and Daryl Gillett (Southern Arava Agricultural Research and Development Center), together with Noach Natanel (Arava Institute) for monitoring the field experiment in Yotveta. We would like to thank Dr Heike Gnad and Dr Jon Falk (Saaten Union, Gatersleben, Germany) for their professional and accurate working of the doubled haploids from interspecific crosses outside their regular protocols. Special thanks to the Fridman lab members for their valuable work in the field trials and in the postharvest. This work was supported by the US-Israel Binational Agricultural Research and Development Fund (BARD) (grant#IS-5658-24C) to E.F and D.K., by the CAPITALISE/Horizon2020 (grant#MD-862201-2) to E.F, and a National Science Foundation CAREER grant (IOS-2046256) to D.K.

## Author Contribution

E.F. made the initial design of the CMPP with E.B. who performed the F_1_, BC_1_ and BC_2_ crosses; S.B. organized the phenotypic and genotypic datasets, designed and performed the analyses, and wrote the initial draft for the manuscript; L.D.T. performed the chloroplast DNA extraction and sequencing; A.B. managed the seeds for the field trials; S.Y. provided genotyping data for the training of the GWAS models and was involved in the final writing of the manuscript; R.P.A. generated the DNA libraries for the WGS; D.K. performed the WGS bioinformatics including variant calling, imputation, filtering and was involved in writing the manuscript.

